# Identification of common molecular biomarker signatures in blood and brain of Alzheimer’s disease

**DOI:** 10.1101/482828

**Authors:** Md. Rezanur Rahman, Tania Islam, Md. Shahjaman, Julian M.W. Quinn, R. M. Damian Holsinger, Mohammad Ali Moni

## Abstract

**Background:** Alzheimers disease (AD) is a progressive neurodegenerative disease characterized by memory loss and confusion. Neuroimaging and cerebrospinal fluid-based early detection is limited in sensitivity and specificity as well as by cost. Therefore, detecting AD from blood cell analysis could improve early diagnosis and treatment of the disease. The present study aimed to identify blood cell transcripts that reflect brain expression levels of factors linked to AD progression.

**Methods:** We analyzed blood cell and brain microarray gene expression datasets from NCBI-GEO for AD association and expression in blood and brain. We also used eQTL and epigenetics data to identify AD-related genes that were regulated similarly in blood and brain.

**Results:** We identified 9 differentially expressed genes (DEG; AD versus controls) common to blood cells and brain (CNBD1, SUCLG2-AS1, CCDC65, PDE4D, MTMR1, C3, SLC6A15, LINC01806, and FRG1JP) and 18 genes (HSD17B1, GAS5, RPS5, VKORC1, GLE1, WDR1, RPL12, MORN1, RAD52, SDR39U1, NPHP4, MT1E, SORD, LINC00638, MCM3AP-AS1, GSDMD, RPS9, and GNL2) that were commonly dysregulated between AD blood and brain tissues using SNP and cis-eQTL data. This data revealed significant neurodegeneration-associated molecular pathways in the ribosomal and complement systems. Integration of these different analyses revealed dysregulation of hub transcription factors (SREBF2, NR1H2, NR1H3, PRDM1, XBP1) and microRNAs (miR-518e, miR-518a-3p, miR-518b, miR-518c, miR-518d-3p and miR-518f) in AD. Several significant histone modification sites in DEGs were also identified.

**Conclusion:** We have identified new putative links between pathological processes in brain and transcripts in blood cells in AD subjects that may enable the use of blood to diagnose and monitor AD onset and progression.

## 1. Introduction

Alzheimer’s disease (AD) is a progressive neurodegenerative disease common among elderly individuals that results in progressively severe cognitive impairment. In the USA, 5.7 million people are currently living with Alzheimer’s and this is expected to rise to 14 million by 2050 [1]. AD is diagnosed by the presence of extracellular amyloid plaques and intracellular neurofibrillary tangles in the brain and reflect the pathobiological processes that underlies the disease [2]. Although the pathogenesis of AD is multifactorial in nature, the application of molecular methods to improve diagnosis and assessment of AD has yet to provide substantiated results and hence the quest for early AD biomarkers in peripheral blood has received increased attention [3, 4]. Successful identification of such blood molecular biomarkers will have a high impact on AD diagnosis, care and treatment[5, 6].

Positron emission tomography (PET) based neuroimaging techniques and cerebrospinal fluid are both used in clinical practice to diagnose Alzheimer’s [7, 8, 9]. However, these procedures suffer serious limitations, including the invasiveness of collecting CSF as well as the sensitivity, specificity, cost and limited access of neuroimaging [10]. Considering the shortcomings of available resources for the detection of neurodegenerative diseases, many studies have attempted to explore biomarkers in the blood of AD patients [11, 12]. Circulating cells and proteins are easily accessible from fresh blood samples, the collection procedure is less invasive. Since central mechanisms underlying progression of the disease is still not clear, much attention has been drawn to systems biology approaches as a new avenue to elucidate the possible roles of biomolecules in complex diseases such as AD [13, 14, 15, 16]. For example, evidence of involvement of miRNA deregulation in the development of neurodegenerative diseases has emerged [17, 18]. Consequently, biomolecules such as mRNAs, transcription factors (TFs), miRNAs (and mRNA gene transcripts targeted by such TFs and miRNAs) are increasingly being scrutinised for use as new biomarkers for AD. In addition, the role of epigenetic modifications is also a focus of much interest, with evidence for their importance in the development and progression of diseases such as AD [19, 20]. DNA methylation and histone modifications are common mechanisms for epigenetic regulation of gene expression [21]. It is well understood that factors such as lifestyle, age, environment and co-morbid states effect epigenetic changes [21] as well as risk of AD and that gene methylation and histone modification may be implicated as mediators [21].

We employed an integrative approach to identify molecular biomarker signatures that are expressed under similar genetic control in blood cells and brain in AD using transcriptome and expressed quantitative loci (cis-eQTL). Gene over-representation analysis was performed on core DEGs followed by gene ontology (GO) analysis. Pathway analysis was then used to enrich the DEGs. Core DEGs were further analyzed to identify regulatory factors (TFs, miRNAs) that may affect the DEG in AD-affected tissues, as well as analysis to identify histone modification sites within the identified DEGs. This study specifically focused on biomarker signatures at both transcriptional (mRNAs and miRNAs) and translational levels (hub proteins and TFs) as our intention was to present valuable information that would clarify mechanisms in AD that may provide efficacious potential biomarkers for early diagnosis and systems medicines (Figure 1).

**Figure 1:**
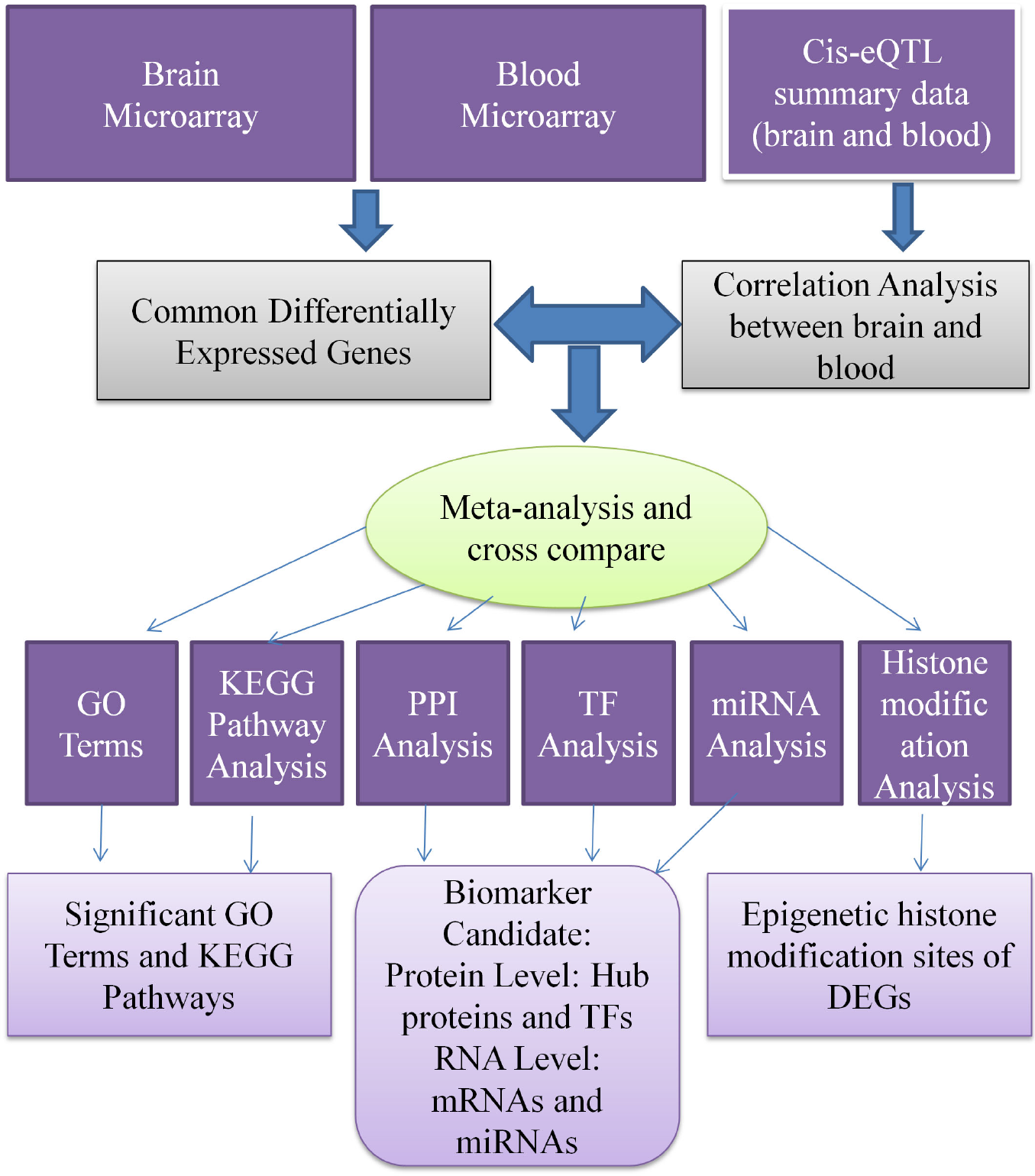
The systems biology pipeline employed in this study. Gene expression datasets of blood of AD were obtained from the GEO and GTEx portal. The datasets were analyzed using in bioconductor environment in R to identify common DEGs between brain and blood tissue. The significantly enriched pathways, GO terms were identified through enrichment analyses. PPI network was analyzed to identify hub proteins. TF-target gene interactions and miRNA-target genes interactions was studied to identify regulatory biomolecules.

## 2. Materials and Methods

### 2.1. Identification of Differentially Expressed Genes from High-throughput Microarray Datasets

We obtained two gene expression microarray datasets GSE18309 (peripheral blood tissues) and GSE4757 (brain tissue) of AD patients from NCBI-GEO database [22]. We then applied a logarithmic transformation to both blood and brain microarray datasets to approximate the datasets to normality and to mitigate the effect of outliers. Following this we applied linear Models for Microarray (limma) through the bioconductor platform in R in order to identify the DEGs from each dataset. The overlapping DEGs between the two datasets were considered for further analysis. We then screened for statistically significant DEGs that satisfied an adjusted p-value ¡0.05 and absolute values of log2 fold for control ¿=1.0. The Benjamini-Hochberg (BH) method was used to adjust p-values.

### 2.2. Geneset Enrichment Analyses to Identify Gene Ontology and Molecular Pathways

We performed geneset enrichment analysis via Enrichr [23] to identify GO and pathways of the overlapping DEGs. The ontology comprised of three categories: biological process, molecular function and cellular component. The p-value¡0.05 was considered as the cut-off criterion for all enrichment analyses.

### 2.3. Protein-protein Interaction Network Analysis

We retrieved the PPI networks based on the physical interaction of the proteins of DEGs using STRING database [24]. A confidence score of 900 was selected in the STRING Interactome. Network visualization and topological analyses were performed through Network-Analyst [25]. Using topological parameters, the degree (greater than equal 18 degree) were used to identify highly interacting hub proteins from PPI analysis.

### 2.4. Identification of Histone Modification Sites)

Histone modification data for the hub genes were retrieved from human histone modification database [26].

### 2.5. Identification of Transcriptional and Post-transcriptional Regulators of the Differentially Expressed Genes

We used TF-target gene interactions from TRANSFAC [27][25] and JASPAR databases [28] to identify TFs. The miRNA-target gene interactions were obtained from miRTarBase [29]. We identified significant miRNAs and TFs (p¡0.05) via Enrcihr [23].

### 2.6. eQTL Effects Between Blood and Brain Tissues

We used eQTL data of both blood and brain from the GTEx Portal which is a database for Genetic Association data (https://gtexportal.org/home/). These eQTL databases link gene SNPs to gene expression. We used them to identify genes with similar genetic control of expression in the two tissues using meta-analysis approaches.

If we allow *x̃* to be the estimated effect of the top-linked cis-eQTL for a gene, we can calculate *x̃* based on the method explained in [30] and as below:

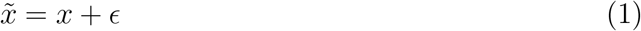

where *x* is the true effect and is the estimated error. The co-variance of the estimated cis-eQTL effects between tissues *i* and *j* across genes can be partitioned into the co-variance of true cis-eQTL effects and the co-variance of estimation errors. Thus We can estimate the correlation of true cis-eQTL effect sizes across genes between tissues *i* and *j*.

### 2.7. Cross Validation of the Candidate Biomarkers

We cross checked the identified common DEGs using AlzGene database, which is a collection of published AD genetic association studies [31]. We also cross compared the identified miRNAs with blood based miRNA signatures and detected 12 diagnostic miRNAs [32].

## 3. Results

### 3.1. Identification of Common Deferentially Expressed Genes between Blood and Brain Tissues

We analyzed microarray gene expression datasets of brain and blood. The analysis revealed 9 (nine) common DEGs (CNBD1, SUCLG2-AS1, CCDC65, PDE4D, MTMR1, C3, SLC6A15, LINC01806, and FRG1JP) in blood and brain.

### 3.2. Identification of AD-associated Genes in Blood that Mirror Those in Brain from eQTL

We used a meta-analysis approach to identify genes from GTEx database that display a similar expression pattern in both blood and brain tissues using eQTL database that link gene variants (SNPs) to gene expression. Thus, we identified 673 blood-brain co-expressed genes (BBCG) using the correlation and meta-analysis approach as explained in the methods section. We identified 18 genes (HSD17B1, GAS5, RPS5, VKORC1, GLE1, WDR1, RPL12, MORN1, RAD52, SDR39U1, NPHP4, MT1E, SORD, LINC00638, MCM3AP-AS1, GSDMD, RPS9, and GNL2) that were commonly dysregulated between AD blood and brain compared to control tissues using SNP and cis-eQTL data of curated, gold-benchmarked OMIM and GWAS catalogues.

### 3.3. Gene Ontology and Pathway Analysis

To clarify the biological significance of the identified DEGs, we performed a geneset enrichment analysis. The significant GO terms were enriched in biological processes, molecular functions and cellular components (Table 1). The pathways analysis revealed significant differences in the Ribosome, Alternative Complement Pathway, Classical Complement Pathway, Lectin Induced Complement Pathway and Cytoplasmic Ribosomal Proteins (Table 2).

**Table 1:**
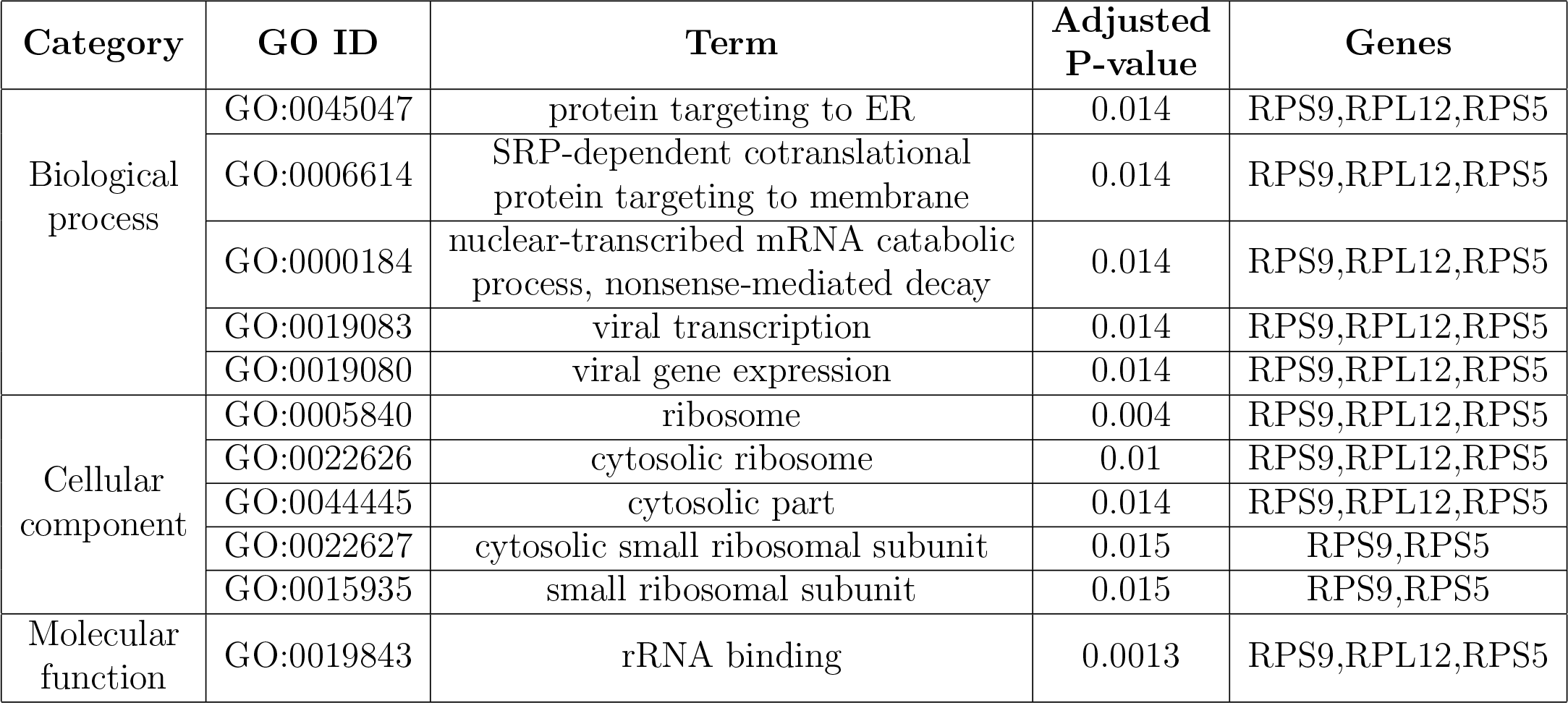
Gene Ontology (biological process, Cellular component and molecular functions) of differentially expressed genes commonto blood cells and brain tissue of Alzheimers disease.

**Table 2:** The significant molecular pathways of common differentially expressed genes between blood and brain tissue of Alzheimers disease.

| Category | Pathways | Adjusted p-values | Genes |
| --- | --- | --- | --- |
| KEGG | Ribosome | 0.021 | RPS9,RPL12,RPS5 |
| BioCarta | Alternative Complement Pathway | 0.020 | C3 |
|  | Classical Complement Pathway | 0.020 | C3 |
|  | Lectin Induced Complement Pathway | 0.020 | C3 |
| WikiPathways | Cytoplasmic Ribosomal Proteins | 0.005 | RPS9,RPL12,RPS5 |

### 3.4. Protein-protein Interaction Analysis to Identify Hub Proteins

A protein-protein interaction network was constructed, encoded by the DEGs to reveal the central protein, the so called hub proteins considering the degree measures (Figure 2). RPS5, RPL12, RPS9, GNL2, PDE4D, and WDR1 were identified as the hub proteins. These are potential biomarkers and may lead to new AD therapeutic targets.

**Figure 2:**
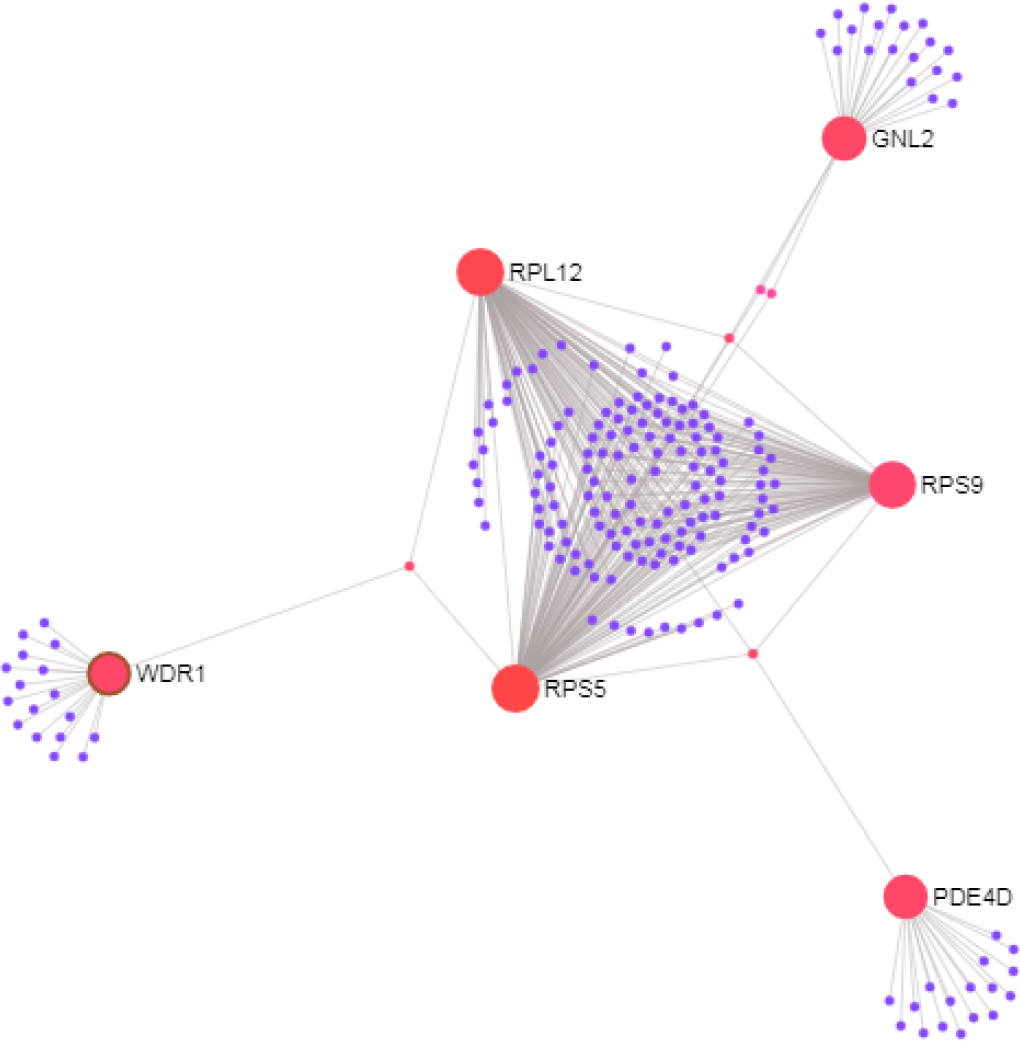
Protein-protein interaction network of the differentially expressed genes (DEGs) in Alzheimers disease. The nodes indicate the DEGs and the edges indicate the interactions between two genes.

### 3.5. Epigenetic Regulation of the Differentially Expressed Genes

In order to identify the probable epigenetic regulation of the hub genes, histone modification data for six eight hub genes (Table 3) were retrieved from HHMD. Table 3 shows that all the hub genes were associated with several histone modification sites.

**Table 3:** Histone modification patterns (obtained from HHMD) of hub genes with respect to the already known histone modification sites in neurodegenerative diseases.

| Official Symbol of DEGs<br>and Hub Genes | RefSeq ID | Histone modification sites already known<br>in neurodegenerative diseases |  |  |  |
| --- | --- | --- | --- | --- | --- |
|  |  | H3K27 | H3K4 | H3K9 | H4R3 |
| RPS5 | NM_001009 | ✓ | ✓ | ✓ | ✓ |
| PDE4D | NM_001104631 | ✓ | ✓ | ✓ | ✓ |
| RPL12 | NM_000976 | ✓ | ✓ | ✓ | ✓ |
| RPS9 | NM_001013 | ✓ | ✓ | ✓ | ✓ |
| GNL2 | NM_013285 | ✓ | ✓ | ✓ | ✓ |
| WDR1 | NM_017491 | ✓ | ✓ | ✓ | ✓ |

### 3.6. Identification of Post-transcriptional Regulator

We identified TFs and miRNAs and their targeted DEGs to reveal regulatory biomolecules that may regulate the expression of DEGs at transcriptional and post transcriptional levels (Table 4). The analysis revealed significant TFs (SREBF2, NR1H2, NR1H3, PRDM1, and XBP1) and miRNAs (miR-518e, miR-518a-3p, miR-518b, miR-518c, miR-518d-3p, and miR-518f) (Table 5) played significant roles in the regulation of the DEGs identified this study.

**Table 4:** Transcription factors with a significant regulatory role of differentially expressed genes that are common to blood and brain tissue in Alzheimers disease.

| **Term** | **Genes** | **P-value** |
| --- | --- | --- |
| SREBF2 | GSDMD,C3,WDR1,HSD17B1,RPL12,NPHP4 | 0.008 |
| NR1H2 | GLE1,GSDMD,SORD | 0.02 |
| NR1H3 | GSDMD,VKORC1,GLE1,SORD,GNL2 | 0.03 |
| PRDM1 | MTMR1,WDR1,PDE4D,SLC6A15,MT1E | 0.04 |
| XBP1 | RPS9,PDE4D,SORD | 0.04 |

**Table 5:**
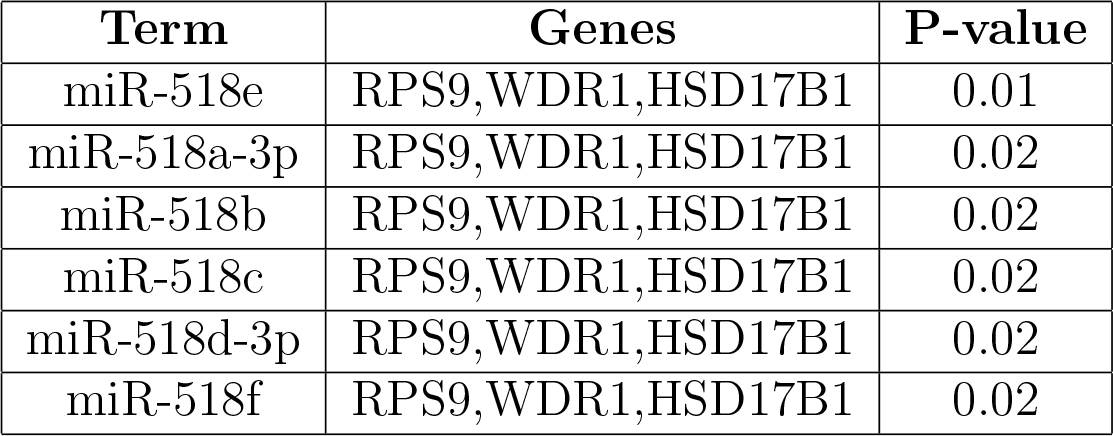
MicroRNAs that significantly regulate common differentially expressed genes between blood and brain tissue in Alzheimers disease.

### 3.7. Cross-validation of the Candidate Biomarker Biomolecules

The hub protein genes were compared with the 618 AD-related biomarkers deposited in the AlzGene database. The result revealed no overlap of these DEGs with AD-related biomarkers in the AlzGene database. We then compared the identified miRNAs with aberrantly regulated blood miRNAs that were reported in a study that detected 12 diagnostic miRNAs of AD patients from peripheral blood samples. No overlapping miRNAs were identified.

## 4. Discussion

The lack of peripheral blood biomarkers for AD has led to a race to identify much needed evidence for the early diagnosis this debilitating disease. The identification of peripheral biomarkers may also shed light on molecular mechanisms of AD and enable the monitoring of treatment. Advances in biomedical technology have spurred discoveries in numerous research areas. Microarray analysis is widely used in biomedical research and is considered a main resource for candidate biomarkers. Microarray databases contain a wealth of untapped genomic information. We analyzed two gene expression datasets from peripheral blood and brain of the AD patients in an attempt to identify potential biomarker candidates.

Our analysis revealed 9 DEGs common to the two transcriptomic datasets of blood and brain tissues. Geneset enrichment analyses also revealed AD-associated molecular signalling pathways that included the ribosome and complement systems. Employing protein-protein interaction networks we also identified dysregulated central hub proteins that control many cellular processes. These hub proteins are considered key drivers in the mechanisms underlying the disease [33]. Therefore, we reconstructed the protein interaction network focusing on the DEGs in an attempt to identify related hub proteins. Such proteins have the potential to contribute to the formation and progression of AD. Of the DEGs we identified, mRNA levels of RPS5, a ribosomal protein, has been shown to be increased in the frontal cortex of AD subjects and AD transgenic mice [34].

Epigenetic alterations are present in different tissues during ageing as well as in neurodegenerative disorders such as AD. Epigenetic factors affect lifespan and longevity. AD-related genes exhibit epigenetic changes, indicating that epigenetics might contribute to pathogenic changes observed in dementia. Epigenetic modifications are reversible and may potentially be targeted by pharmacological intervention. Epigenetic drugs may be useful for treatment of major health problems [35]. We have identified epigenetic changes in hub genes (Tables 3) and have investigated histone modification patterns of DEGs. Histone modifications are posttranslational modifications of the amino-terminal tails of histone proteins that affect nucleosome structures and gene accessibility to TFs. Histone modification thus affects downstream molecular interactions, thereby affecting patterns of gene expression. We report several histone modification sites present within these DEGs and hub genes (Table 3), many of which are already known to be associated with several neurodegenerative diseases [36]. The identification of these known modifications in known genes further validates the discovery of the novel DEGs and hub genes that we have identified in this investigation.

Our analysis also revealed differentially expressed DEGs, TFs and miRNAs that strongly influence gene expression at the transcriptional and post-transcriptional levels (Table 4 and Table 5).

SREBF2: The SREBF2 is a cholesterol regulating genes and significant increased mRNA levels expressesion were observed in the late onset AD in brain and blood microarray observations suggesting SREBF as biomarkers of AD at pathogolical and gene expression levels Picard et al. [37]. In another study evaluated the SREBF2 mRNA level expression in neurodegenerative prion disease. Significant increased expression of SREBF2 was in prion infected neuron cells suggesting cholesterogenic upregulation as neuroalrespone to proion infection emphasizing cholesterol biosynthesi a crtical pathways in prion disease [38]. NR1H3: The genetic variant was studied to determine the effects of rs7120118 variation in the NR1H3 gene on the progression of AD. Significant increase in the mRNA levels of NRIH3 among the AD patients was found by qPCR analysis. Overall, these data suggest that the CT genotype of rs7120118 associates with increased mRNA levels of NR1H3, but the disease severity does not affect NR1H3 expression [39]. Additionally, association analysis of common variants in NR1H3 identified rs2279238 conferring a 1.35-fold increased risk of developing progressive MS. Protein expression analysis revealed that mutant NR1H3 alters gene expression profiles, suggesting a disruption in transcriptional regulation as one of the mechanisms underlying MS pathogenesis. Novel medications based on NR1H3 models are expected to provide symptomatic relief and halt disease progression by reducing the inflammatory response and promoting remyelination [40]. NR1H2: The genetic polymorphism in NR1H2 may contribute to the pathogenesis of AD [41]. PRDM1: The exome sequencing and functional studies revealed the genetic variants of PRDM1 in Crohn’s [42].PRDM1 was associated with systemic lupus erythematosus(SLE)[43]. There is a link between Cerebral inflammation and degeneration in systemic lupus erythematosus [44], but inverse relations suggested for SLE and parkinsons disease patients since SLE had a decreased risk of subsequent Parkinson disease [45]. However, study indicates that the risk of dementia may be elevated in individuals with SLE, an autoimmune disease affecting a range of systems including the peripheral and central nervous system concluding SLE is significantly associated with dementia [46]. WDR1: WDR1 is associated with adaptive immunity highlighting its central role immunologic synapses [47] and cardiovascular diseases [48][47]. GNL2: GNL2 plays role in neurogenesis of retina in Zebrafish[49]

RPS9 and RPS12: Gene Ontology (GO) annotations related to gene RPS9 and RPS12 include structural constituent of ribosome according to Genecards summary, but the role of these ribosomal proteins in the neurodegenerative disease is obscure.

XBP1: The role of XBP1 in neurodegeneration remains controversial and appears to be disease-specific. XBP1 occupancy was observed on the promoters of genes linked to neurodegenerative pathologies including Alzheimers disease [50], although the relevance of these events remains speculative. Indeed, XBP1 activates a plethora of target genes involved in a variety of physiological functions, including neuronal plasticity [51][50][52], suggesting an important role during the branching and maturation of developing neurons. Accumulation of unfolded or misfolded proteins in the ER leads to an ER stress response, which is characteristic of cells with a high level of secretory activity and is implicated in a variety of disease conditions such as AD [53]. Hub protein PDE4D was particularly noteworthy since recent studies have suggested that phosphosdiesterases are promising therapeutic drug targets in AD [54].

miRNAs play important roles in gene regulation and there is emerging evidence demonstrating their potential for use as biomarkers for AD and other diseases; it is likely therefore that miRNAs play significant roles in the pathogenic process underlying AD [55][56]. Indeed, such roles have been suggested for miR-518e and miR-518a-3p in AD [51]. Similarly, miR-518c may also be a useful biomarker for Parkinson’s disease [57] while miR-518b is dysregulated in esophageal carcinoma [58].

**Table 6:**
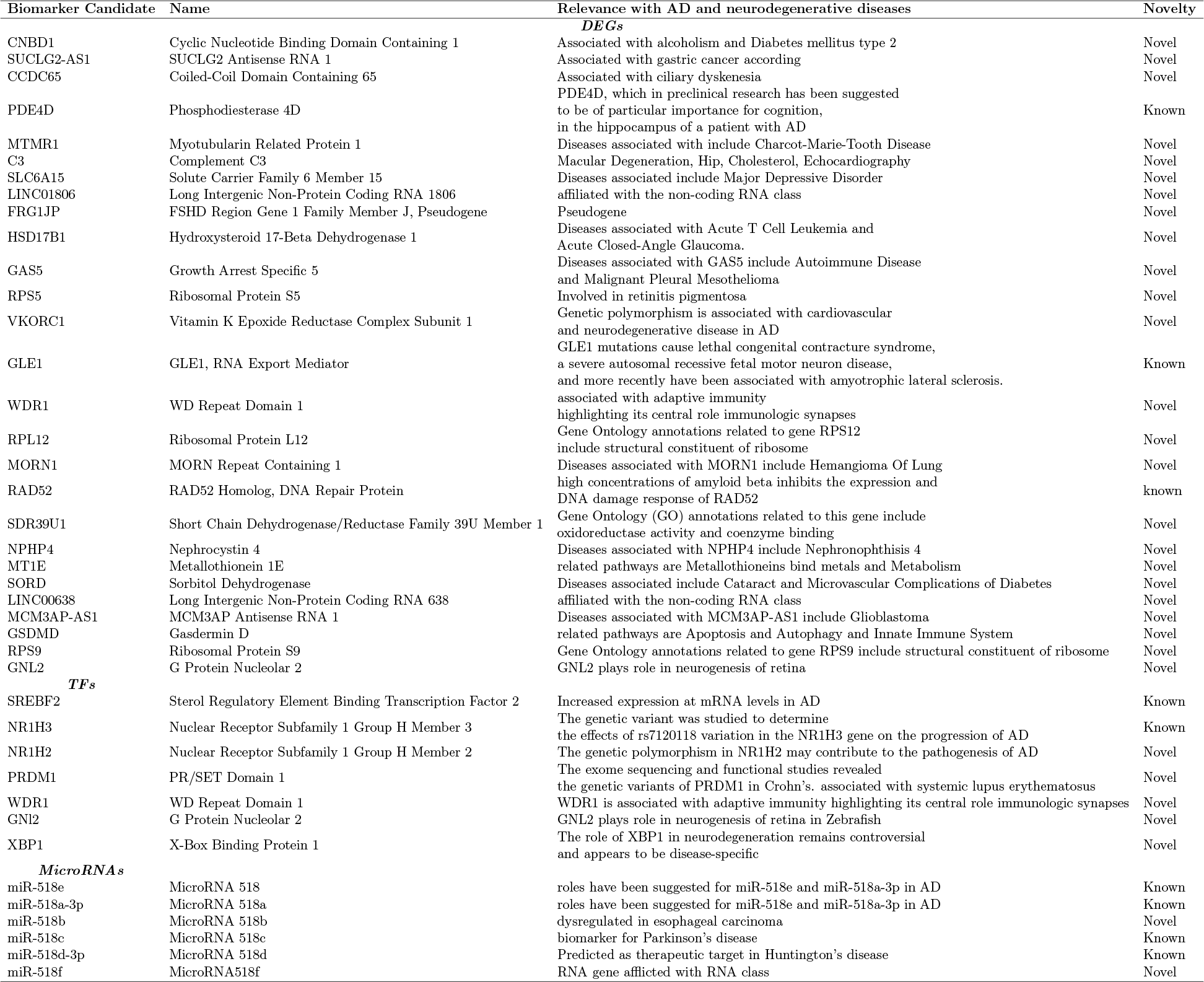
A List of Biomarker Candidates Proposed in the Present Study for AD.

## 5. Conclusion

In the present study, we analyzed blood and brain transcriptomic and eQTL data to identify common DEGs between these two tissues in Alzheimer’s disease. We integrated these common DEGs into pathway analysis for protein-protein interactions, TFs and miRNAs. Nine common DEGs were identified from microarray data of blood and brain. We also identified 18 eQTL genes common to blood cells and brain cells. Neurodegeneration-associated molecular signalling pathways and several miRNAs were identified as putative transcriptional and post-transcriptional regulators of the DEGs we identified. In addition, several histone modification sites of hub proteins were were also identified. Thus, we have identified potential biomarker transcripts that are commonly dysregulated in both blood cells and brain tissues. We propose that these biomarkers may enable the rapid and cost effective assessment of blood sample analysis for the diagnosis of AD. This novel approach to identify markers can be employed in easily accessible tissue (blood) to assess its expression in an inaccessible tissue (brain), and is one that could be applied to other related clinical problems. We now propose a more detailed validation of this approach and of the putative biomarker transcripts we have identified with clinical-based investigations.

